# Efficacy of Mating Disruption Treatments Against Spongy Moth (Lymantria dispar dispar) Applied Using Unmanned Aerial Vehicles

**DOI:** 10.64898/2025.12.10.693540

**Authors:** Ksenia S. Onufrieva, Andrea D. Hickman, Tom W. Coleman

## Abstract

Unmanned aerial vehicles (UAVs) are increasingly used in precision pest management, yet their performance in operational forest settings remains underexplored. We evaluated the efficacy of SPLAT® GM-Organic mating disruptant applied using an UAV at the dosages of 14.8 and 11.4 g AI/ha for control of the spongy moth, *Lymantria dispar dispar* L. (Lepidoptera: Erebidae). Both treatments reduced trap catches by >90% for 10 weeks following the application, meeting the efficacy requirement set by the USDA’s National Slow the Spread (STS) Program. One year after the application, trap catches continued to be reduced by 28% and 67% in plots treated with 14.8 and 11.4 g AI/ha, respectively. These levels of trap catch reduction in the year of treatment and one year after the treatment application are comparable to the ones observed following fixed-wing aerial treatments. These results indicate that UAV-applied SPLAT® GM-Organic meets STS requirements for operational use and is suitable for integration into the program for treating small or isolated blocks. These findings also have broader implications for the use of unmanned aerial vehicles to deploy SPLAT® formulations in forest pest management programs.

## Introduction

Unmanned aerial vehicles (UAVs) are increasingly being used in pest management, but their performance under operational conditions, particularly in forested systems, remains underexplored. Combining a UAV with a precise application payload is expected to provide a safer, more economical method of aerial treatment application with minimized off-target movement of the sprayed agent (1). Piloted aircraft are limited by high operational complexity and cost (2), are unsuitable for small and/or fragmented areas as well as areas bordering water sources (3) and are shown to fail in small treatment blocks due to inadequate pheromone coverage (4). As a result, pest management programs are frequently unable to treat all areas in need of protection, while private growers and homeowners may resort to conventional insecticides that are potentially harmful to non-target organisms. Integrating UAV technology into pest management programs offers a promising solution by enabling more targeted applications and expanding treatment access to previously inaccessible areas. However, without clear guidance and field-based validation, the potential of UAVs may remain unrealized. As stated in the USDA’s National Roadmap for Integrated Pest Management (2018), effective pest control requires the use of “available technology to prevent unacceptable levels of pest damage by the most economical means, while posing the least possible risk to people, property, resources, and the environment.” UAV applications can help achieve that goal only if their use is systematically tested, refined, and integrated into operational programs.

Many countries outside the United States have long used UAVs in agricultural applications; for example, Japan relies on UAVs for 90% of all aerial applications for field crops (5). In the U.S., recent field trials have begun to build a foundation for broader adoption by demonstrating that UAV-based pesticide applications can achieve efficacy comparable to that of conventional aerial applications (6) and ground-based sprayers (7). In cotton production, Cavalaris et al. (8) found that UAV applications of defoliants were more efficient than both ground and aerially applied systems, highlighting the potential for UAVs to improve operational flexibility and reduce overall chemical use. Trials in cranberry systems have demonstrated that UAV-applied pheromones can significantly reduce trap catches and mating success of two Lepidopteran pests (9). Collectively, these studies support the growing role of UAVs as effective delivery platforms for pest management in agriculture.

However, empirical evaluations of UAV applications remain limited in forested environments. The present study addresses this gap by evaluating the efficacy and persistence of UAV-applied SPLAT® GM-Organic (ISCA, Riverside, CA) for mating disruption of the spongy moth, *Lymantria dispar dispar* L. (Lepidoptera: Erebidae). Spongy moth is one of the most destructive invasive insects in North American forests. It is a federally regulated pest and managed through three strategies: suppression, eradication, and the National Slow the Spread (STS) Program (10). STS is an area-wide integrated pest management program implemented at the leading edge of the expanding population front with the goal to reduce the rate of spongy moth spread across the United States. In the STS Program, mating disruption is the primary method of spongy moth control, and since 2017, SPLAT® GM-Organic has been used exclusively on more than 100,000 ha annually, with an average treatment block size of 2,029 ± 113 ha (11).

Additionally, given that SPLAT® is a biologically inert matrix used in a range of semiochemical-based formulations designed for controlled release of an active ingredient (12-17), the results of this study may provide insights for other SPLAT-based management programs. Finally, while UAVs are often presented as a technological breakthrough in precision agriculture and pest management, some authors have cautioned against uncritical adoption of such narratives driven by media hype and inflated expectations (18). Field-based trials – including this one – are therefore essential to evaluate the actual effectiveness of UAV-based treatments under operational field conditions, which can represent fluctuations in topography, isolated forest stands, limited sight lines, and varying forest canopy heights. Furthermore, the STS Program requires new methods for applying mating disruption treatments and the ability to treat small (∼200 ha or smaller), isolated populations.

## Methods

### 1. Efficacy of the mating disruption treatment in the year of application

Eight experimental plots, each 100 m x 100 m, were established in Goshen Wildlife Management Area, VA, within forest stands dominated primarily by oak (*Quercus* spp.) that had experienced minimal human disturbance or active forest management for over five years. Plots were established in either a valley bottom or along the edge of hill, but all contained varying elevational changes (Fig. 1). Four plots were left untreated and used as a control, and four plots were treated on May 26, 2023, with SPLAT® GM Organic using an agricultural drone sprayer M6E-X (Beijing TT Aviation Technology Ltd, Beijing, China) equipped with an Aksata (formerly Agridrones, Kfar Saba, Israel) Ointment Application System (OAS) precise applicator payload (Fig. 1). The experimental plots were spaced at least 700 m apart from each other to prevent treatment interference (19). Three plots were treated with an overall dosage of 14.8 g/ha (6 g AI/acre), which is an operational dosage in the STS Program. One plot was treated with an overall dosage of 11.4 g AI/ha (4.6 g AI/acre) due to a calibration error. The drone flew at a speed of 10 m/sec 1.2 m (4 feet) above canopy treating plots at an average rate of 36 seconds per 0.4 ha (one acre). SPLAT® GM-Organic was dispensed from a 250 g-capacity cartridge compatible with a manual caulking gun. Each cartridge was weighed before and after application to determine the amount of product applied per plot. The product was extruded in a continuous line by steady pressure from the tube, with 15 m spacing between adjacent lines (Fig. 2). During application, the material intermittently broke into large, cohesive masses that were deposited on the target surfaces (Fig. 3).

**Figure 1.**
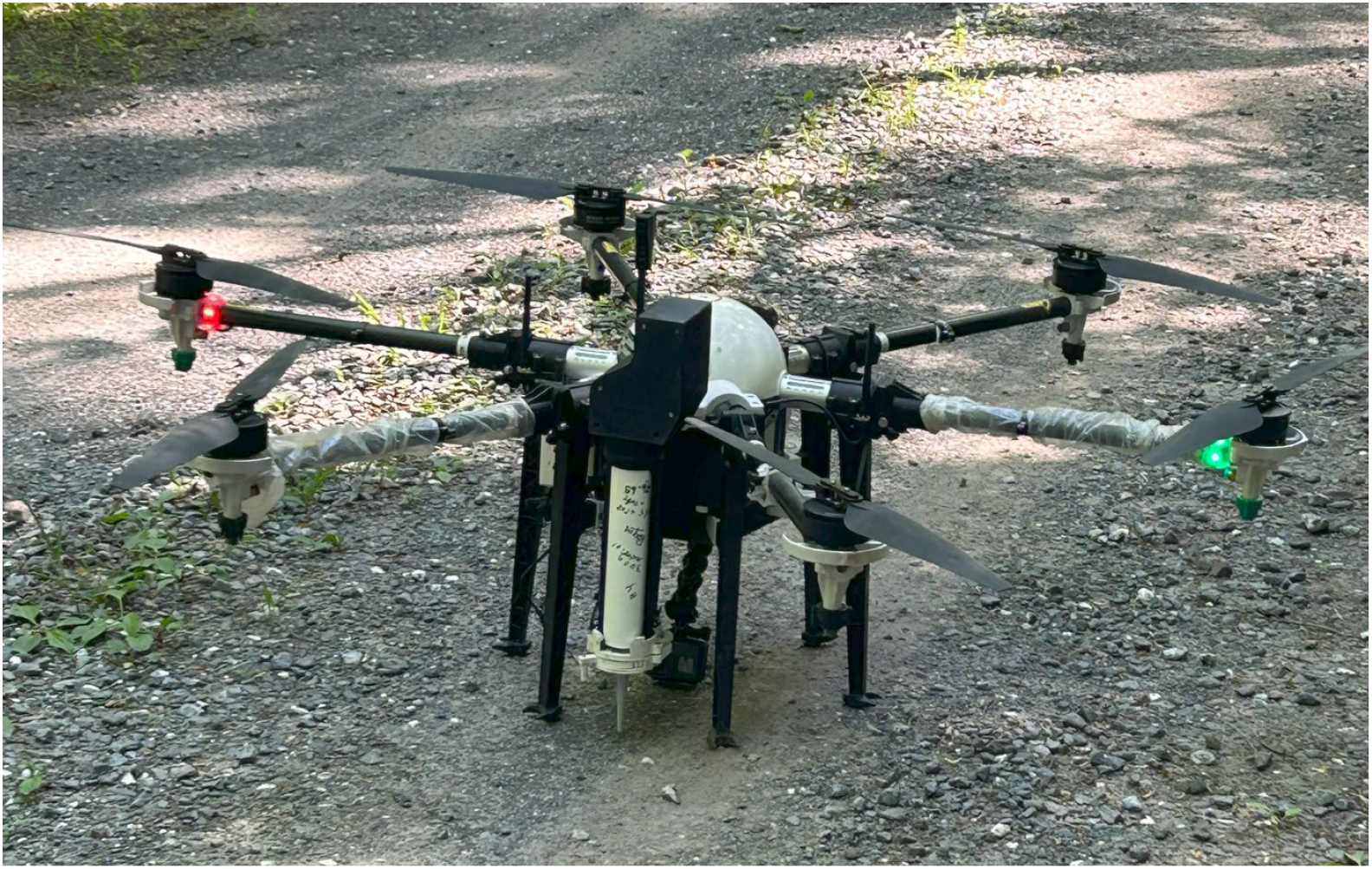
Unmanned Aerial Vehicle [Agricultural drone sprayer M6E-X (Beijing TT Aviation Technology Ltd)] equipped with Aksata (formerly Agridrones) Ointment Application System (white vertical tube) to apply SPLAT® GM Organic.

**Figure 2.**
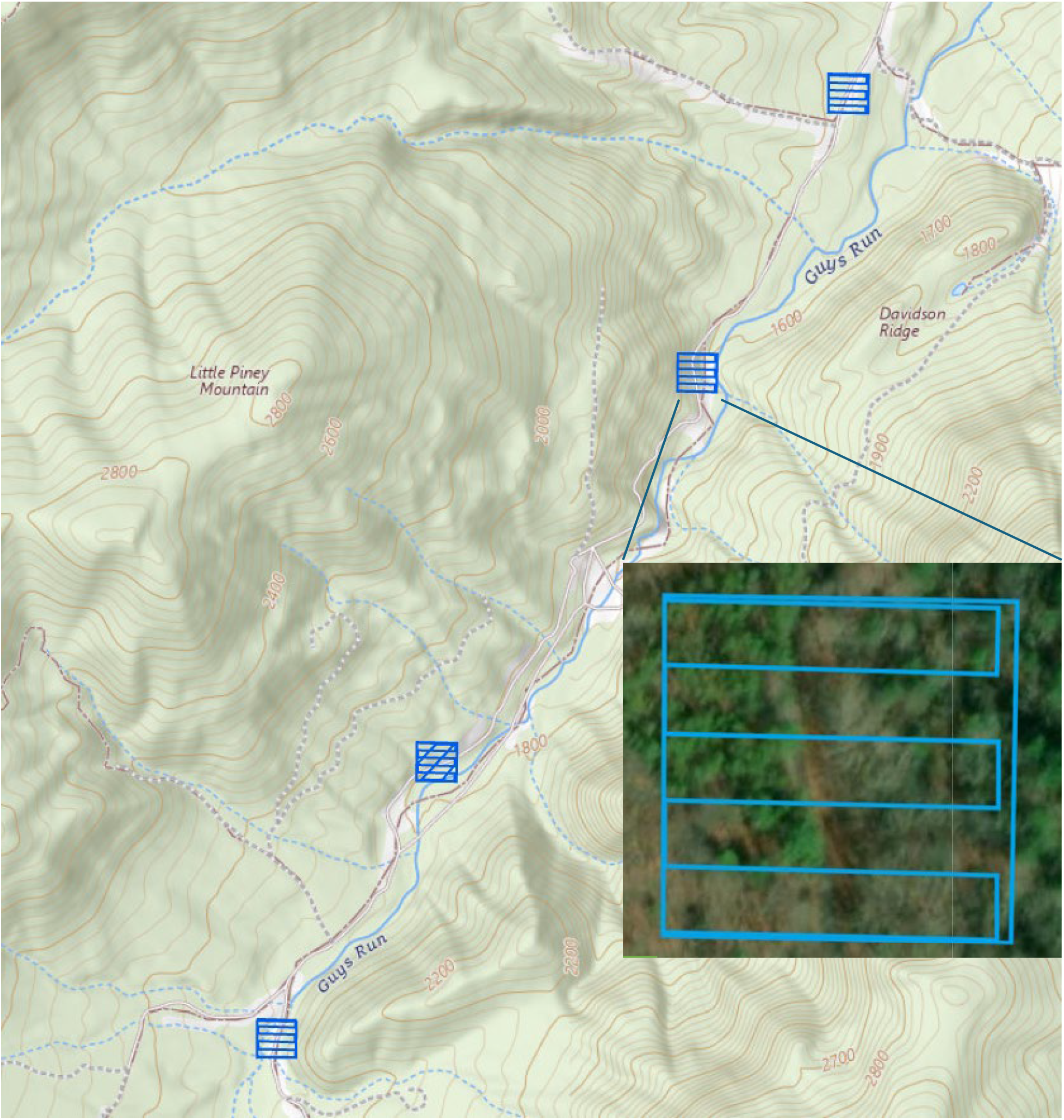
Experimental plots treated using the Agricultural drone sprayer M6E-X (Beijing TT Aviation Technology Ltd) equipped with Aksata Ointment Application System to apply SPLAT® GM Organic. Flightlines are shown in black on the main map and in blue on the insert.

**Figure 3.**
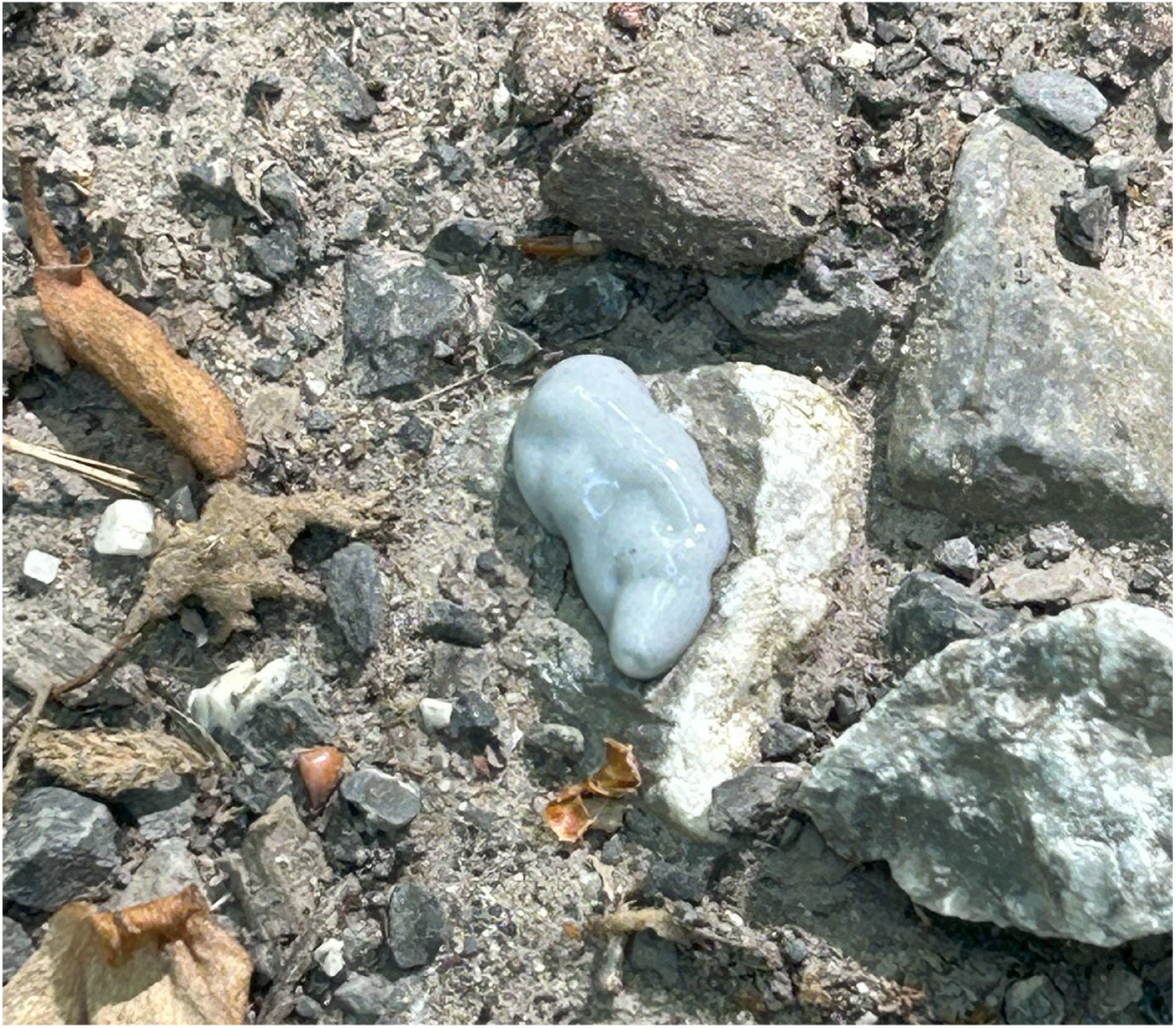
Large, cohesive masses of SPLAT® GM-Organic deposited by UAV on target surfaces. The experimental plots were monitored from week two to week ten after the treatment application (June 6 – August 4, 2023), week 1 was skipped due to logistic reasons. We assessed the efficacy of the applied pheromone treatments according to our standard protocol for the formulation efficacy test using lab-reared adult males (20, 21). In each plot, we established a male moth release point surrounded by four USDA milk carton pheromone-baited traps (Fig. 4).

**Figure 4.**
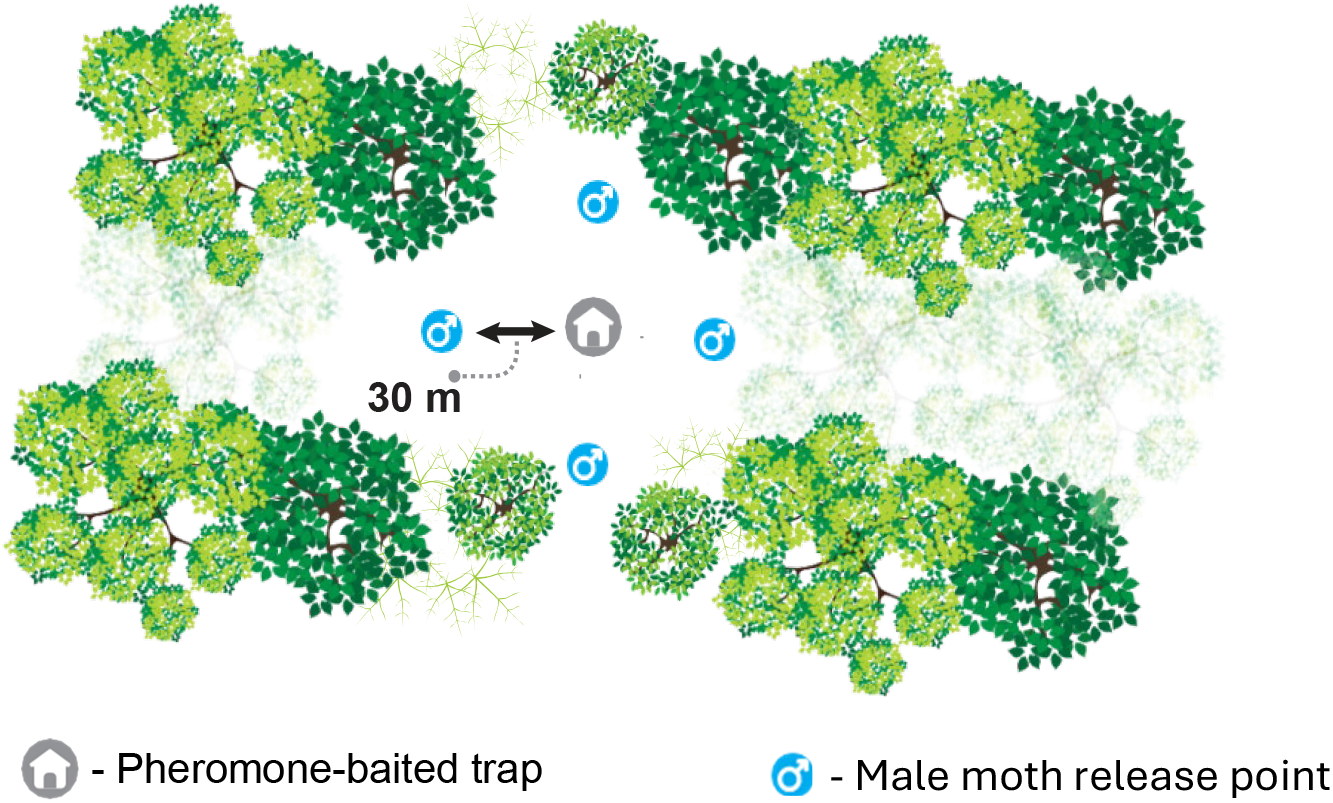
Experimental plot design established to determine the efficacy of mating disruption treatments (Figure courtesy of Michael Stamper, Virginia Tech).

We released an equal number of spongy moth males per release (50 - 100) 2-3 times per week for a total of ten releases. This spongy moth release rate adequately simulates population densities currently controlled by mating disruption treatments (22). Mating success was evaluated by monitoring male moth recaptures in pheromone-baited traps. Only lab-reared males were used for the analysis to ensure equal population densities among experimental plots. Marked spongy moth males were supplied by the USDA APHIS, Pest Survey Detection and Exclusion Laboratory (Buzzards Bay, MA), and reared to adults in the field lab just prior to release. A solvent red 26 dye (Royce International, Paterson, NJ) was added to the larval diet at the rearing facility. The dye is expressed in adults and is used to differentiate between released and feral male moths.

### 2. Efficacy of the mating disruption treatment one year after the application

In 2024, we monitored the experimental plots treated in 2023 with SPLAT® GM-Organic at 14.8 g AI/ha and used the same protocol to evaluate the effect of that treatment on spongy moth mating success one year after the application. In the untreated control plots and the plots treated in 2023, we established a male moth release point surrounded by four pheromone-baited traps and released equal number of males per release (50 – 100). We made six male moth releases between June 14 and July 31, 2024. Mating success was evaluated by monitoring male moth recaptures in pheromone-baited traps. Only lab-reared males were used for the analysis to ensure equal population densities among experimental plots.

### 3. Statistical analysis

We used a linear mixed-effects model (mixed model ANOVA) to compare trap catches between control and treated plots. Treatment and week were treated as fixed effects, plot was included as a random effect to account for repeated measures and variation among plots, and plot-by treatment interaction used as an error term. Model fitting was performed using restricted maximum likelihood (REML) (JMP Pro 18, 2025, SAS Institute Inc.). Post-hoc comparisons among treatments were conducted using Tukey’s Honestly Significant Difference (HSD) test. Least squares means and standard errors were extracted for pairwise comparisons.

To evaluate overall and weekly percent trap catch reductions, we calculated percent reduction relative to the untreated control using raw means for each treatment group, following Abbott’s formula (23). We used the STS ≤10% of control trap-catch threshold (24) as a descriptive reference; figures display raw means ± SE to facilitate visual comparison with program-defined efficacy thresholds.

## Results and Discussion

### 1. Efficacy of the mating disruption treatment in the year of application

Trap catches in the pheromone-treated plots were significantly reduced compared to the untreated control plots (*F*_2,57_ = 599, *P* = 0.003; Fig. 5). Across all treated plots, season-long trap catch was reduced by more than 90% relative to the untreated controls. In plots treated with 14.8 g AI/ha (operational dosage), total trap catch was reduced by an average of 97%, ranging from 95% to 99% among plots. Notably, the plot treated with 11.4 g AI/ha (due to a calibration error) also showed a 99% reduction in trap catch.

**Figure 5.**
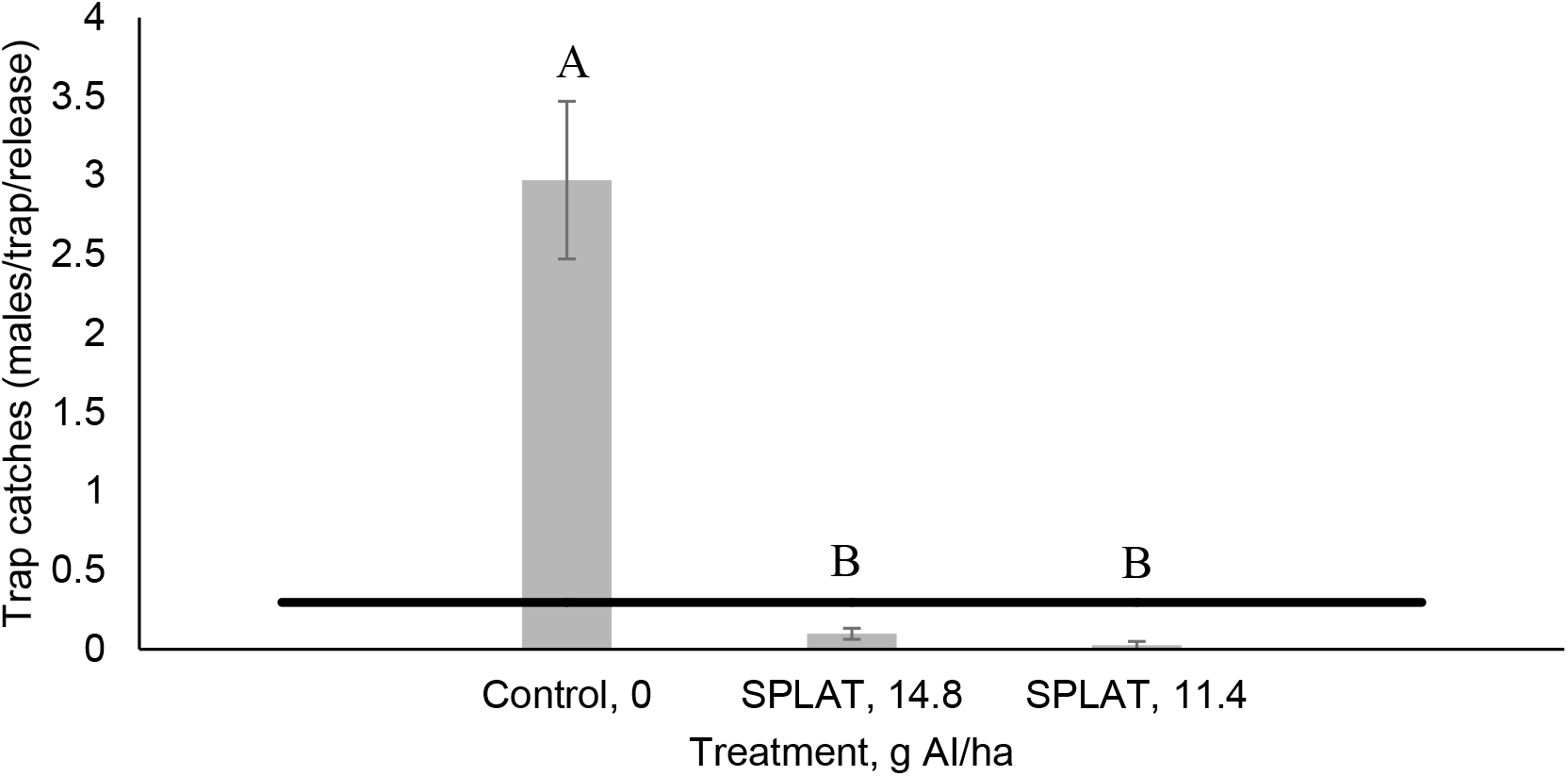
Mean trap catches (± SE) recorded in pheromone-treated and untreated control plots. The black horizontal line represents the 10% mating success threshold used in the STS Program; bars below this line indicate successful treatment. Bars with the same letter are not significantly different (Tukey’s HSD: P > 0.05).

In the STS Program, treatment is considered effective if it results in ≥90% reduction in trap catch for at least eight weeks post-application (24) to cover the entire period of spongy moth flight (up to six weeks) and to provide a safety margin for uncertainties associated with the logistics of treatment planning and with songy moth phenology. Therefore, overall reduction in trap catch does not necessarily indicate that treatment efficacy was maintained throughout the entire adult flight period. To evaluate the consistency of treatment efficacy, we analyzed weekly trap catch data. In all treated plots, weekly trap catch was reduced by more than 95% for ten consecutive weeks (Fig. 6), confirming that the treatment maintained full efficacy throughout the season.

**Figure 6.**
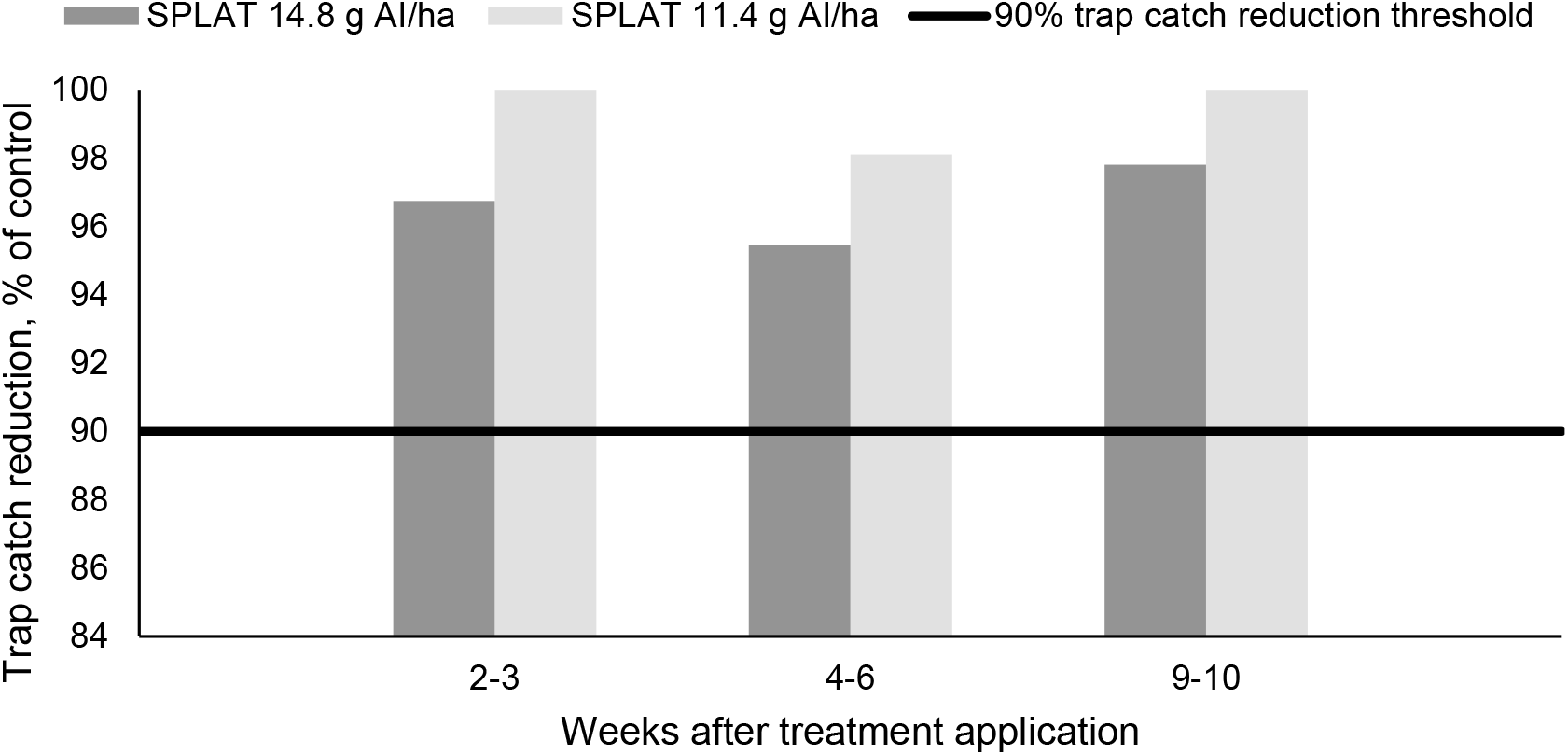
Weekly percent trap catch reduction in plots treated with SPLAT® GM-Organic at 14.8 g AI/ha (dark grey) and 11.4 g AI/ha (light grey). The black horizontal line represents the 90% trap catch reduction threshold used in the STS Program; bars above this line indicate successful treatment.

### 2. Efficacy of the mating disruption treatment one year after the application

In the STS Program, persistence of pheromone treatments is an operational concern. While long-lasting pheromone residues may provide some additional suppression of target populations, they can also interfere with trap-based monitoring (25). Spongy moth pheromone is known to persist in the environment for a year or more, potentially complicating post-treatment evaluations (21).

In 2024, one year after the UAV application, season-long trap catches in plots treated with 14.8 g AI/ha and 11.4 g AI/ha were reduced by 28% and 67%, respectively, relative to the untreated control plots. This trap catch reduction one year after the treatment application is comparable to the trap catch reductions observed one year following fixed-wing aerial treatments (26). Although season-long trap catch reduction varied among the treated plots (2 – 47% in plots treated with 14.8g AI/ha dosage and 67% in the plot treated with 11.4g AI/ha), these differences were not statistically significant (Fig. 7).

**Figure 7.**
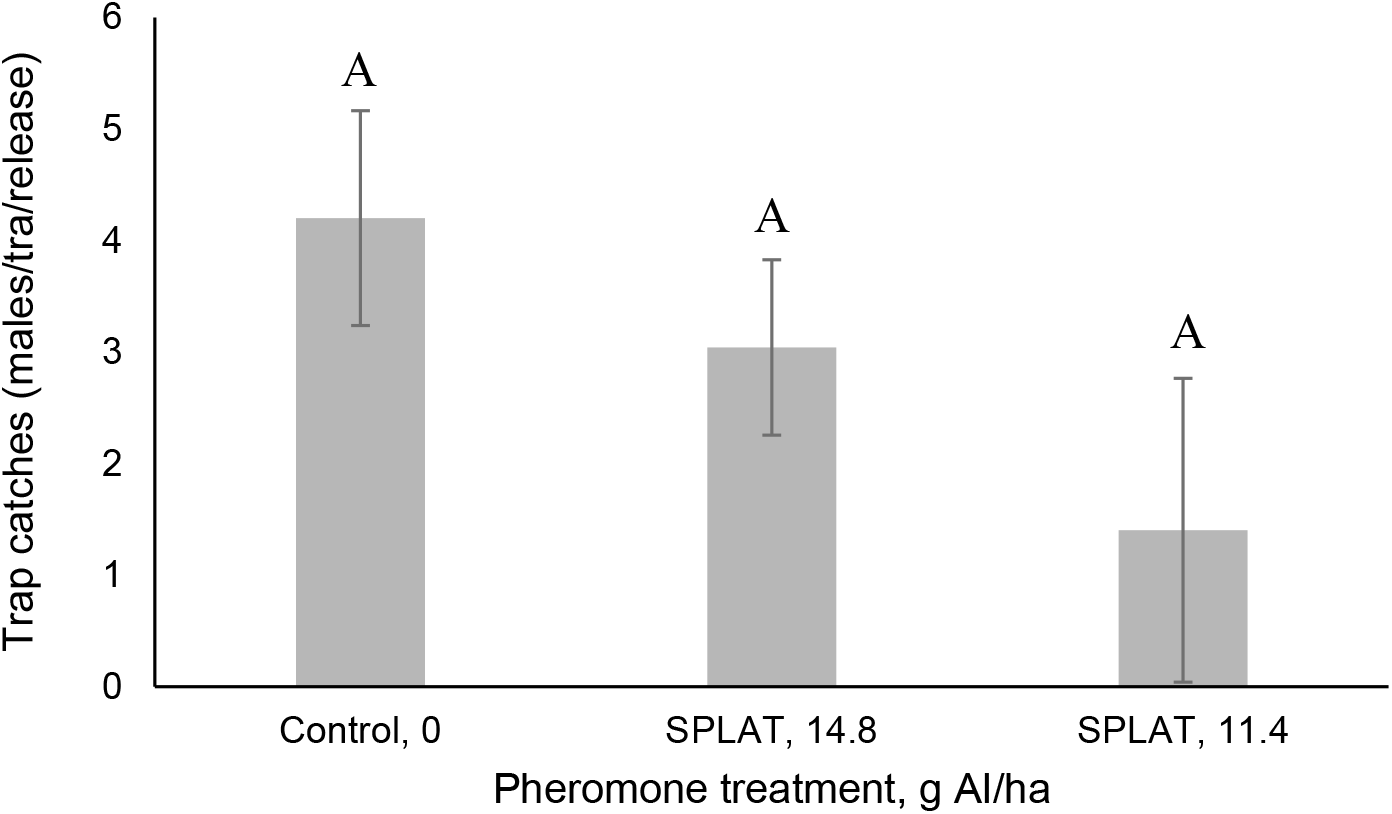
Mean trap catches (± SE) recorded in treated and untreated control plots. Bars sharing the same letter are not significantly different (Tukey’s HSD: P > 0.05).

Previous work has shown that droplet size distribution can strongly influence the persistence of SPLAT® GM-Organic, with larger droplets (≥3100 µm) releasing pheromone over a longer period compared to smaller droplets (Onufrieva et al. 2025, in review). In the present study, UAV application produced significantly larger cohesive masses resulting in fewer individual deposits compared with conventional aerial applications (Fig. 3), which would be expected to result in stronger second-year suppression. However, the modest reductions observed here suggest that droplet size alone may not be a reliable predictor of long-term efficacy. Other factors, such as weathering, canopy interception, or application pattern, may limit persistence even when deposits are large. From an operational perspective, the fact that UAV-applied treatments yielded a second-year effect similar to conventional fixed-wing applications means that post-treatment evaluation protocols can remain consistent regardless of application method.

Taken together, these results indicate that UAV-applied mating disruption treatments meet STS Program efficacy thresholds, reducing trap catch by over 90% for at least ten weeks. In addition, the observed reduction in trap catches one year after application suggests that UAV-applied treatments produce a similar level of residual effect as conventional aerial treatments. These findings support the inclusion of UAV-based mating disruption into operational spongy moth management programs and suggest that no additional adjustments to post-treatment evaluations are necessary.

In the STS Program, the mean treatment block size for aerially applied mating disruption is 2,029 ± 113 ha (11). In contrast, UAVs are typically deployed in much smaller blocks, with the capacity to treat approximately 200 ha (500 acres) per day and a minimum operational block size of about 2 ha (5 acres) (Aksata, 2025, pers. Comm.). Treatment costs for UAV applications range from $20 to $60 per 0.4 ha (one acre). Integrating UAVs with fixed-wing aircraft in an operational program could maximize budget efficiency and enable treatment of all critical areas without leaving untreated gaps or applying excessive coverage.

Many management programs rely on SPLAT® formulations, which are currently applied by hand (14, 15, 17, 27), using ground-based equipment (12), or aerially via fixed-wing aircraft (20). Each of these application methods presents unique operational challenges due to labor, terrain, cost, or logistical constraints. UAV-based delivery could overcome these limitations by enabling faster, more uniform, and more precise applications than manual or ground-based methods, and more cost-effective, targeted, and flexible deployment than fixed-wing aerial systems.

By demonstrating that UAV-applied SPLAT® GM-Organic meets performance standards in both the year of treatment and the following season, this study contributes to the growing body of evidence supporting UAVs as practical tools for semiochemical-based pest management and offers protocols for UAV-applied mating disruption applications.

## Conclusion

UAV-applied SPLAT® GM-Organic meets the requirements for operational use in the National Slow the Spread Program against spongy moth.

## Acknowledgements

We thank Hannah Nadel, Christine McCallum, and Susan Lane (USDA APHIS PPQ) for supplying spongy moth pupae; Jacob Ninio, Ari Eisenberg (Aksata Technologies, Los Angeles, CA), and Tslil Yitzhak (Tslil Yitzhak Ltd., North District, Israel) for providing the aerial applications; ISCA Technologies for providing SPLAT® GM-Organic; Donald Duerr (USDA, Forest Service) for administrative support; Mannin Dodd (Virginia Tech) for technical assistance; Mikeal Jolly (Goshen Scout Reservation) for providing facilities and tremendous support; Ron Hughes, Gene Sours, Bill Mohler, Kent Burtner (VA Department of Game and Inland Fisheries) for providing forest access; Department of Entomology at Virginia Tech for providing facilities; Michael Stamper (Virginia Tech Libraries) for assistance with figure preparation. This work was supported by the USDA Forest Service, Forest Health Protection (Grants number 19-DG-11083150-005 to K.S.O.). Mention of a proprietary product does not constitute an endorsement or a recommendation for its use by USDA.

## Conflicts of Interest

The authors declare no conflict of interest.

## Notes

### Competing Interest Statement

The authors have declared no competing interest.

